# Single-Cell Transcriptome Analysis Identifies Novel Biomarkers Involved in Major Liver Cancer Subtypes

**DOI:** 10.1101/2022.02.01.478756

**Authors:** Asish Kumar Swain, Prashant Pandey, Riddhi Sera, Pankaj Yadav

## Abstract

Liver cancers including hepatocellular carcinoma (HCC) and intrahepatic cholangiocarcinoma (ICC) are leading cause of death worldwide. Single-cell transcriptomics studies have vast potential in advancing our understanding of cancers by defining the cellular composition of different solid tumor types. We peformed an integrated analysis using single-cell RNA sequencing (scRNA-seq) data from cancerous and healthy liver tissues in order to identify the molecular progression and intercellular heterogeneity across cell types in the liver cancer. Moreover, we performed a subtype specific analyses, separately for HCC and ICC, to identify any molecular drivers uniquely associated with these liver cancers. The scRNA-seq dataset comprising 5 healthy controls and 19 liver cancer patients were collected from Human Cell Atlas and Gene Expression Omnibus (GEO), respectively. Our analyses confirmed upregulation of four previously known malignant cell marker genes, namely, EPCAM, KRT19, KRT7 and S100P in the cancerous liver cells. Of these, KRT7 gene has been reported to be associated with ovarian cancer in the past studies. Noteworthy, four marker genes specific to the G1/S (MCM5 and PCNA) and G2/M phases (HMGB2 and CKS2) of the cell cycle were upregulated in the cancerous liver cells. This indicates that these four marker genes are actively dividing in these two phases in cancerous cells as compared to normal liver cells. Our differential expression analysis identified 2 upregulated genes (ATF3 and S100A11) and 2 downregulated genes (FCN3 and FGB) in the liver cancer. Our subtype based differential expression analysis identified 4 genes (HSPA6, LMNA, ATP1B1 and DCXR) specific to HCC and 3 genes (HSPB1, APOC3 and APOA1) specific to ICC. CD4+ T-cell, Hepatocyte, neutrophil, mesenchymal cells and liver bud hepatic cells are the predominant cell-types in liver cells. Our scRNA-seq study revealed the mesenchymal cells as potential malignant cell types in liver cancers. Our work suggests future research on developing liver cancer subtypes therapies could target these cell types and associated molecular markers.

## Introduction

The human liver plays a central role in the metabolism and immune function. Liver cancer (LC) is the most frequent fatal malignancy in the world. The prognosis for liver cancer is poor. Three major types of liver cancers include intrahepatic cholangiocarcinoma (ICC), hepatocellular carcinoma (HCC) and hepatoblastomas[1][2]. Of these, HCC is ranked third leading cause of deaths occurring due to various cancers [3]. Moreover, 90% of all the cases of primary liver cancers belong to the HCC[4]. The ICC is the second most common liver cancer after HCC [5]. The ICC is associated with poor prognosis and very little is known about the involved biological abnormalities[6]. In the past, several studies have been done in order to understand the physiology, development, and pathology of the human liver. Due to diverse microenvironment, genomic variation, immense heterogenity at cellular level and variability in disease progression cause challenge in design therapeutic treatment for liver cancer. To tackle the unsolved problems, understanding of liver tumor at intratumoral (with in tumor), intertumoral (HCC/ICC) is crucial for targeted therapy and new treatment modelling. Recent study using single-cell RNA sequencing (scRNA-seq) data on immune cells from different tissue sites in the liver shows that the nature of the immune cells is dynamic in the cancerous microenvironment[7]. In another study, the inter-tumor diversity of cells of ICC have been studied using scRNA-seq techniques to get insights into the intercellular crosstalk[8].

Single-cell transcriptomics studies have vast potential in advancing our understanding of cancer etiology. High-throughput single-cell RNA sequencing (scRNA-seq) techniques are available at a reasonably low cost[9]. The advent of scRNA-seq has enhanced our ability to build detailed cellular maps of tissues, gaining unique insights into the cancer microenvironment[10]. The scRNA-seq allows the gene expression to be studied at the cellular resolution which makes in-depth characterization of cellular heterogeneity easier. Recent efforts in collecting scRNA-seq data has been quite successful. For instance, the *Human Cell Atlas* project was initiated with objective to classify all cell types present in different organs of the human body based up on the gene expression profile[11].

Despite several studies in the past, a systematic analysis of the cellular composition of the liver tumor microenvironment at subtype level remained unexplored as yet. In particular, an integrated scRNA-seq analysis of liver cancer subtype is required to better understand the diease etiology. In this study, we retrieved 4 different healthy and cancer (*HCC* and *ICC*) scRNA sequencing data from public databases. To identify cellular heterogeneity at subtype level, data integration of healthy and cancer datasets and clustering of similar cell types were performed and visualized. Cell cycle analysis was done to gain insight of proliferation level of distinct cell populations. Furthermore, differential expression analysis at cell cluster and sample level was carried out to identify the top marker genes. Using top marker genes, rare cell type annotation and gene enrichment analysis was performed to understand the biological functions of DEGs. Our study findings in comparative mapping of liver cancer subtypes at cellular and genetic level can be a theoretical basis for new drug designing, targeted therapeutic and help researchers to understand the underlying mechanism of tumor progression and chemotherapy resistance at subtype level.

## Materials and Methods

### scRNA-seq Datasets

To compare the gene expression patterns between the healthy and cancerous liver cells, four different scRNA-seq datasets were obtained from publicly available databases. Our first scRNA-seq dataset (e.g., *Healthy1*) comprised 30,000 liver cells from 5 healthy samples collated at the Human Cell Atlas[11]. The second scRNA-seq dataset (e.g., *LC*) comprised 9,946 liver cells from 19 LC patients available at the GEO database (accession number GSE125449)[12]. The LC patient cohort included 11 males and 8 females in the age range 41 – 80 years. The third scRNA-seq dataset (e.g., *HCC*) was obtained from GEO database which comprised 32154 liver cells from 2 HCC patients (GSE166635)[13]. Our last scRNA-seq dataset comprised 31,302 liver cells from 5 ICC samples (e.g. *ICC)* and 3 adjacent non-tumor samples (e.g. Healthy2) (GSE138709)[8].

### Quality Control

The pre-processing of the four scRNA-seq datasets was performed separately following same steps using the *Seurat* package in R software version 4.0.4. At first, the dead cells and empty barcodes were removed. Next, the low-quality cells having less than 200 genes were removed. Also, genes that are expressed in less than 3 cells were removed. To further filter low-quality cells, we removed the high mitochondrial transcript (50%) containing cells. This high threshold was chosen as normally hepatocyte cells have higher proportion of mitochondrial genes. The doublet (i.e two cells captured in single droplet) filtering of cells was not considered for the reason that potential doublet may be a single large cell with two nuclei[14]. **Table.1**. shows the summary of cells remained in the four datasets after the quality control steps. Next, to remove the impact of potential outlier’s log-normalization was performed in each dataset where gene counts were divided by total counts in the cell then multiplied by scale factor(eg. 10000) and then log transformed the scaled values.

**Table 1.** Summary of 4 datasets [Healthy, Liver cancer (*LC*), HepatoCellular Carcinoma (*HCC*), Intrahepatic Cholangio Carcinoma (*ICC*)] and number of cells and genes remained before and after Quality Control(*QC*)

| Dataset | Database & Accession No | No of samples | Number of Cells & genes before QC |  | Number of Cells and genes after QC |  |
| --- | --- | --- | --- | --- | --- | --- |
| Healthy1 | HCA (Human Cell Atlas), GSE115469 | 5 | 30000 | 37011 | 26998 | 27523 |
| LC | NCBI GEO, GSE125449 | 19 | 9946 | 21324 | 9946 | 21287 |
| HCC | NCBI GEO, GSE166635 | 2 | 32154 | 33694 | 31990 | 22594 |
| ICC | NCBI GEO, GSE138709 | 5 | 17090 | 19408 | 17090 | 19408 |
| Healthy2 | NCBI GEO, GSE138709 | 3 | 16901 | 16559 | 16901 | 16559 |

### scRNA-seq Dataset Integration

The healthy samples (i.e., *Healthy1* dataset) was combined with Cancer dataset (i.e *LC)* and for subtype analysis adjacent non tumor samples(i.e *Healthy2* dataset) combined with subtype cancer samples(i.e. *HCC & ICC)* in order to perform different comparative analyses. For the purpose of data integration, 2000 high variable features (or genes) were extracted separately from the healthy and the three cancer datasets. The *FindIntegrationAnchors* function in the *Seurat* package was used to find the inter sample anchor between healthy and cancer datasets. Next, the data integration step was performed using the *IntegrateData* function available in the *Seurat* package. This function outputs the concatenated centered expression matrix.

### Comparative Analysis of Healthy and Cancer Samples

The comparative analyses were performed between the healthy and cancerous samples using the three integrated datasets (see above). The comparative analyses included the comparison of the number of distinct genes per cell, total count of genes per cell, percentage of mitochondrial genes per cell, and percentage of ribosomal protein genes per cell.

### Cell-types Clustering and Annotation

The pre-processed data was used to perform the clustering of similar cell types. For each integrated datasets, we extracted top 2000 highly variable genes (HVGs) having a high cell-to-cell variation in the expression level. Next, linear dimension reduction was performed on the selected top 2000 HVGs by using principal component analysis (PCA). We used the top 30 principal components (PCs) to generate K-Nearest Neighbor (KNN) and Shared Nearest Neighbor (SNN) graphs. The KNN and SNN clustering was performed using the Louvain algorithm with a resolution parameter of 0.1. For the subtypes analyses (i.e., HCC and ICC integrated datasets), the resolution parameter was set at 0.2 to get the detailed rare cell type variation[15]. Further, we performed dimension reduction using non-linear uniform manifold approximation and projection (UMAP) and t-Distributed stochastic neighbor embedding (t-SNE) using the top 30 PCs[16][17]. Top two components of UMAP and t-SNE were used to visulalise the gene expression variations across cells for LC, HCC and ICC.

To find the top expressed genes (marker genes) from each cluster, Seurat “FindMarkers” function was used where Wilcoxon Rank sum test used to calculate the average log-fold change for each gene. Genes having average log fold change > ± 1.20 and adjusted p-value<0.05 used to identify predominat cell types(cell type annotation) in all clusters.In order to indentify the cell-type corresponding to a given gene marker, we used the *CellMarker* database (URL: *http://biocc.hrbmu.edu.cn/CellMarker/)*.

### Cell-Cycle Analysis

The variation in the gene expression profiles of single cells in different cell-cycle phases can interfere with the functional analysis scRNA-seq data. Therefore, we calculated the cell-cycle phase specific scores based up on the expression level of 97 marker genes previously known to regulate the G1/S (43 marker genes) and G2/M (54 marker genes) phases of cell-cycle[18]. A high cell-cycle score indicates proliferative cells, whereas a low score indicates non-dividing cells. The data normalization was performed using the LogNormalize method with default parameter setting[19]. In this method, first the relative copy number for each gene is calculated in a cell. Then, the relative copy number counts are multiplied by a scaling factor (default value is 10,000). Finally, natural log-transformation was performed. The expression level of genes specific to G1/S and G2/M phases were visualized using the Dotplot.

### Differential Expression Analysis

Top differentially expressed genes (DEGs) for each subtype level (Healthy1∼LC, Healthy2∼HCC, Healthy2∼ICC) was identified by using Wilcoxon’s rank sum test. Up and down regulated genes (p-value <0.05) were visualised using volcano plot. Genes having average log-fold change > ±1.20 at each subtype level considered more significant for liver tumor. Out of them, DEGs specific to HCC and ICC was identified. The expression level of the malignant cell marker genes was analysed in the integrated datasets using t-SNE plot.

### Gene Enrichment Analysis

We performed gene enrichment analysis in order to get insights into the biological function of the top DEGs. For this purpose, we selected top 20 DEGs from the largest cluster in the t-SNE plot using the *Clusterprofiler* package in R[20].

## Results

### Comparative Analysis of Healthy and Cancer Samples

We performed comparative analyses using the three different scRNA-seq datasets separately for LC-integrated, HCC and ICC (**Table 2**). We compared healthy and cancer for total gene count, unique gene count, mitochondrial gene percentage and ribosomal protein gene percentage. **Figure 1** shows the results of comparative analyses for the healthy and three cancer types (LC, HCC and ICC). We removed the high mitochondrial transcript (50%) containing cells. The violin plot shows the mitochondrial gene percentage after preprocessing (**Figure 1E)**.

**Table 2.**
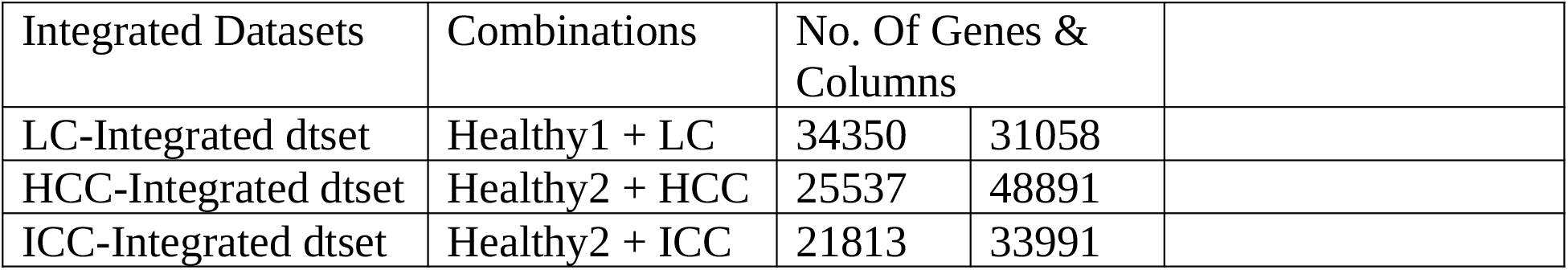
Summary of the 3 Integrated Datasets after pre-processing

**Figure 1.** Violin plot indicating **A) Percentage** of mitochondrial genes i B) Number of distinct gene **C)** Percentage of mitochondrial genes D) Percentage of ribosomal encoding protein genes E) Percentage of mitochondrial genes after processing in both healthy and cancer samples

### Cell-types Clustering and Annotation

We selected top 2000 highly variable genes (HVGs) from each integrated dataset. The top 5 HVGs including IGLC3, HBA2, IGHA1, HBA1 and IGLC2 were found to be common for HCC and ICC (**Figure 2A**). For HCC, the top HVGs included IGHA1, IGKC and, IGHG1 while for ICC the top HVGs were IGLL5, FGB, APOA1 (**Supplementary Figure 5**).

**Figure 2.** Feature selection and dimension reduction A) Top 2000 High variance genes, where top 20 genes labelled B) After dimension reduction scatter plot of expression levels of latent feature PC1 and PC2

**Figure 2B** shows the plot of first two PCs. Top genes associated with the first 4 PCs shown in **Supplementary Figure 3A** and heatmap of first 15 PCs shown in **Supplementary Figure 3B**. The standard deviation of the PCs was plotted shown in **Supplementary Figure 3C**. Based on standard deviation of principal components 30PCs with a larger standard deviation were selected for further analysis. For clustering KNN graph was constructed, based on Euclidian distance in PCA where edge weight between any two cells were refined based on shared overlap in their local neighborhood. Our Cell type clustering identified 12 cell clusters which are composed of healthy and cancerous cells that are visualised using t-SNE and UMAP plots (**Figures 3A and 3B**). By performing subtype anlysis 18 cell clusters obtained from integrated HCC and healthy dataset and for integrated ICC and healthy dataset also 18 distinct cell clusters obtained which are summarised in **Supplementary Figure 7**.

**Figure 3.** Clustering of dimensionally reduced integrated dataset shows **A)** t-SNE plot of integrated dataset **B)** UMAP plot of integrated dataset having 12 distinct clusters **C)**t-SNE plot of the healthy and cancer cells by their sample type **D)** tSNE plot of integrated dataset where cells are grouped by their cell status.

Markers genes for each cluster identified from the integrated dataset, top markers genes from each cluster summaried in **Supplementary Table 1**. By annotating markers genes to the cell markers database among diverse cell population CD4+ Tcell, Hepatocyte, endothelial cell, Neutrophil, Progenitor cell and mesenchymal cells are the top celltypes. Normal and Cancerous liver cells were overlapped and summarised in **Figure 3C**. Some clusters comprise of mostly the cancerous liver cells (Cluster 4,7,9,10,11). Bar plot of percentage of healthy and cancer cells in clusters shown in **Supplementary Figure 6**. suggests Cluster 4 and 7 (Progeniter and mesenchymal cell types) have higher abundance of cancerous cells.

### Cell cycle Analysis

Based on the expression level of G2/M and S phage markers, each cell is assigned a cell score that predict whether the cell is in G2/M or G1 phase. Violin and dot plots of cell cycle phase-specific score summarised in **Supplementary Figures 1A and 1B**, where G1/S phase score found higher in the cancerous cells compared to healthy cells. **Figure 3D**. shows t-SNE plot of cell cycle status of each cell by grouping the cells by cell cycle phase. Histogram of cell count in different cell cycle of each cell type shown in **Figure.4.A. Figure.4.B**. shows expression levels of G1/S and G2/M cell cycle gene in t-SNE plot, cluster 9 shows high expression of G1/S and G2/M cell cycle specific genes which indicate cells in this cluster are actively dividing.

**Figure 4.** **A) Histogram** of cell count in different cell cycle of 12 cell clusters B) Expression level of G1/2 and G2/M cell cycle gene in t-SNE plot **C)**Expression level of G1/S marker genes in healthy and cancer condition **D) Expression** level of G2/M marker genes in healthy and cancer condition

The expression level of marker genes specific to G1/S and G2/M phase is shown in **Figure 4.C and 4.D**, interestingly expression level of phase-specific marker genes high in cancerous liver cells. Genes like PCNA (G1/S marker gene) and HMGB2, CKS2(G2/M marker gene) have overexpression in cancer condition.

### Differential expression and gene enrichment analysis

Top 5 differentially up and down regulated gene between tumor and normal liver sample shown in Table.3. Volcano plot of differential expressed gene in healthy and cancer liver shown in **Figure.5** DEGs with average log2 fold change > **±**1.20 in liver cancer cell shown in **Supp. Table.2**. The expression level of the top marker gene of each cluster is shown in the t-SNE plot (**Figure 6A**), heatmap of selected top differentially expressed gene from each cluster is shown in **Figure 6B**. By subtype analysis, on HCC integrated dataset HSPA6, ATPIB1 genes are upregulated and PPA2, DCXR genes are downregulated. On ICC integrated dataset HSPB1, keratin family genes genes are upregulated and APOC3, APOA1 genes are downregulated whole list of Differential expressed genes in HCC and ICC integrated dataser summarised in **Supp.Table 3 and 4**. Differentially expressed genes of the integrated dataset those are express differentially in HCC and ICC integrated dataset are summarised in **Figure.6.C**.

**Table 3.** Top 5 up and downregulated genes between healthy and cancer samples where avg**_log2FC** shows log fold-change of the average expression between the two groups, **pct.1** explains the percentage of cells where the feature is detected in the first group, **pct.2** explains The percentage of cells where the feature is detected in the second group, **p_val_adjusted** explains Adjusted p-value, based on Bonferroni correction using all features in the dataset.

|  | **avg_log2FC** | **pct.1** | **pct.2** | **p_val_adj** | **Regulation** |
| --- | --- | --- | --- | --- | --- |
| MTRNR2L8 | 5.075821745 | 0.495 | 0.002 | 0 | Up |
| MTRNR2L12 | 4.791436538 | 0.488 | 0.005 | 0 | Up |
| MTRNR2L1 | 4.494704085 | 0.338 | 0.002 | 0 | Up |
| IGHG3 | 4.477730201 | 0.235 | 0.047 | 0 | Up |
| GNB2L1 | 3.992506494 | 0.928 | 0 | 0 | Up |
| MT-TE | -4.988556497 | 0 | 0.585 | 0 | Down |
| MT-RNR1 | -5.587421301 | 0 | 0.929 | 0 | Down |
| SAA2 | -5.749242427 | 0.024 | 0.834 | 0 | Down |
| SAA1 | -6.19806912 | 0.045 | 0.952 | 0 | Down |
| MT-RNR2 | -7.095887133 | 0 | 0.99 | 0 | Down |

Abbreviations: avg\_log2FC = average log fold change, pct.1 = percentage of cells in group1, pct.2 = percentage of cells in group2, p\_val\_adj = Adjusted p-value

**Figure 5.** Volcano plot showing the differentially expressed genes in liver cancer cells as compared to normal liver. In the differential gene expression analysis the most significant genes are those genes which are present on the uppermost section in the plot, i.e., genes which have a high value of –log10P. The genes present on the right side of the plot indicate genes that are upregulated in liver cancer. The genes present on the left side of the plot indicate those genes that are downregulated in liver cancer compared to the normal liver. The genes coloured in blue colour in the centre of the plot indicate the uniformly expressed genes in both the conditions.

**Figure 6.** Differentially expressed genes in each cluster. **A)** t-SNE plot showing one differentially expressed genes from each of the 12 different clusters. **B)** Heatmap showing differentially expressed genes from the 12 different clusters. **C)** Venn diagram of significant genes that are differentially expressed specific to HCC and ICC.

The expression level of 3 marker genes namely, EPCAM, KRT19, and KRT7 in the normal and cancerous cells is shown in **Figure7A**. Interestingly, these 3 genes were underexpressed in almost all the normal cell types, while overexpressed in a few cell types(cluster-4) of the cancerous liver. This suggests that these 3 genes might be associated with liver tumor and could be considered as markers of malignant cells as reported in previous studies. Furthermore, S100P gene was found to be overexpressed in one of the clusters of the cancerous liver while it was underexpressed in the normal liver (**Figure 7B**).

**Figure 7.** Differential Expression analysis. **A)** Expression level of EPCAM, KRT19, and KRT7 in the normal and cancerous liver cells. The upregulation of these genes can be seen in the cells of the cancerous liver. **B)** The expression level of S100P gene in the normal and cancerous liver cells.

We performed gene enrichment analysis using the top 20 differentially expressed genes from the biggest cell type cluster as shown in **Figure 3B**. Molecular function enrichment showed that 6 genes were involved in amide binding and 5 genes were involved in peptide binding functions (**Supplementary Figure 4a)**. The biological process enrichment results showed that 10 genes were associated with neutrophil degranulation, neutrophil activation, neutrophil-mediated immunity, and neutrophil activation involved in immune response (**Supplementary Figure 4b**). The cellular component enrichment results are shown in **Supplementary Figure 4c (**need to mention details as done for molecular function and biological function).

## Discussion

Effective development of therapies against liver cancer warrants deeper understanding of the micro-environment and the gene expression pattern of the cancerous liver. The scRNA-seq has become a powerful high-throughput technique in the last decades with high-resolution for sequencing of the transcriptome. These advancements have facilitated the study of cell-to-cell variation to better understand the complex cellular microenvironment of cancerous tissues[20]. In the present study, we performed a comparative analysis using single-cell transcriptomic data from healthy and cancerous liver tissues.Our results showed that the cancerous liver cells were more actively dividing than the normal liver cells, as expected. Our study validated the upregulation of four previously known malignant cells marker genes, namely, EPCAM, KRT19, KRT7, and S100P in the liver cancer cells. Among these, epithelial cell adhesion molecule (EPCAM)gene is previously known to be highly overexpressed in malignant cells and liver cancer[8]. Moreover, it is reported that EPCAM expression is more pronounced in tumor-initiating cells of many carcinomas. Therefore, EPCAM has been used as a molecular marker to target these tumor-initiating cells[21]. KRT19 is a member of the keratin family, and its upregulation in liver cancer is unknown yet. Thus, understanding the mechanism of action of KRT19 gene can be helpful in patient-specific treatment of liver cancer[22]. Previous studies have shown the upregulation of KRT7 gene in ovarian cancer. However, its expression is highly variable in different types of cancer [23]. In present study, KRT7 was found to be upregulated in the liver cancer. S100P gene is associated with intracellular as well as extracellular funtions andis involved in cell signalling by binding with the receptor for advanced glycation end products[24]. The overexpression of S100P in early stages of different cancers is reported in the past studies. Thus, these four malignant marker genes can act as a molecular target for cancerous liver cells by gene inactivation or blocking their functional pathway[25]–[28].

Further, our differential expression analysis identified the genes which are upregulated and downregulated in the cancer liver cells. The top upregulated genes included S100A11, ATF3 while downregulated genes included FCN3, FGB. By subtype analysis, on HCC integrated dataset HSPA6,LMNA, ATPIB1 genes are upregulated and PPA2, DCXR genes are downregulated. On ICC integrated dataset HSPB1, keratin family genes genes are upregulated and APOC3, APOA1 genes are downregulated Recently it was reported overexpression of S100A11 in mice liver accelerate lipid deposition[29], which may accelerate liver cancer. By upregulation of CYR61 expression ATF3 inhibits tumorigenesis so upregulation of ATF3 may impact on the mechanism[30]. FGB gene which is associated with Dysfibrinogenemia sometimes it may cause cirrhosis and liver tumor, Recently it was found FCN3 can be a prognostic biomarker for immune response in liver cancer, still more research need to be done to validate FCN3.

By subtyping analysis, to validate subtype biomarkers in HCC it was found up regulation of HSPA6 (heat shock proteins) associate with hepatocellular carcinoma and LMNA gene due to proliferation and migration ability function as an oncogene in HCC[31]. PPA2 which modulates histone methylation that interferes with p-53 induced apoptosis can be a marker gene for HCC. Study shows down regulations of DCXR indicates poor prognosis in HCC patients[32].

To validate the biomarkers in ICC, Heat shock protein beta1(HSPB1) which is a negative regulator of ferroptic cancer cell death, upregulation of HSPB1 can be a signification factor for ICC[33]. inhibition of APOC3 which is already used for the treatment of hypertriglyceridemia[34], which associated with excess accumulation of cholesterol in liver can be a biomarker for ICC patients. APOA1 which is already validated as novel serum biomarer for prognosis of HCC, the impact of APOA1 in ICC need to be study further[35]. As Keratin family genes(KRT19, KRT18, KRT8) Overexpressed in ICC samples, some keratin genes (KRT19, KRT7) studied regarding liver cancer, still more study needs to be done.

In conclusion, our work identified novel differentially expressed genes that can act as the potential targets for liver cancer. These findings will help in getting more insights into the liver cancer etiology. Further functional studies can be conducted to understand how these genes interact with each other in the tumor microenvironment. Our work suggests that developing therapies against liver cancer should consider potential role of these identified markers while their mode of action in liver cancer microenvironment is required to be established through experimental validation.

## Acknowledgements

This project was supported by the seed grant XXX funded by Indian Institute of Technology, Jodhpur. The HCA is acknowledged for providing the publicly accessible datasets.

## Author Contributions

AS, PP, RS performed data analyses; PY designed and supervised the study; AS, PP and PY wrote the first manuscript draft; all authors contributed by comments and approved the final manuscript.

## Additional Information

The online version of this article contains a data supplement.

---

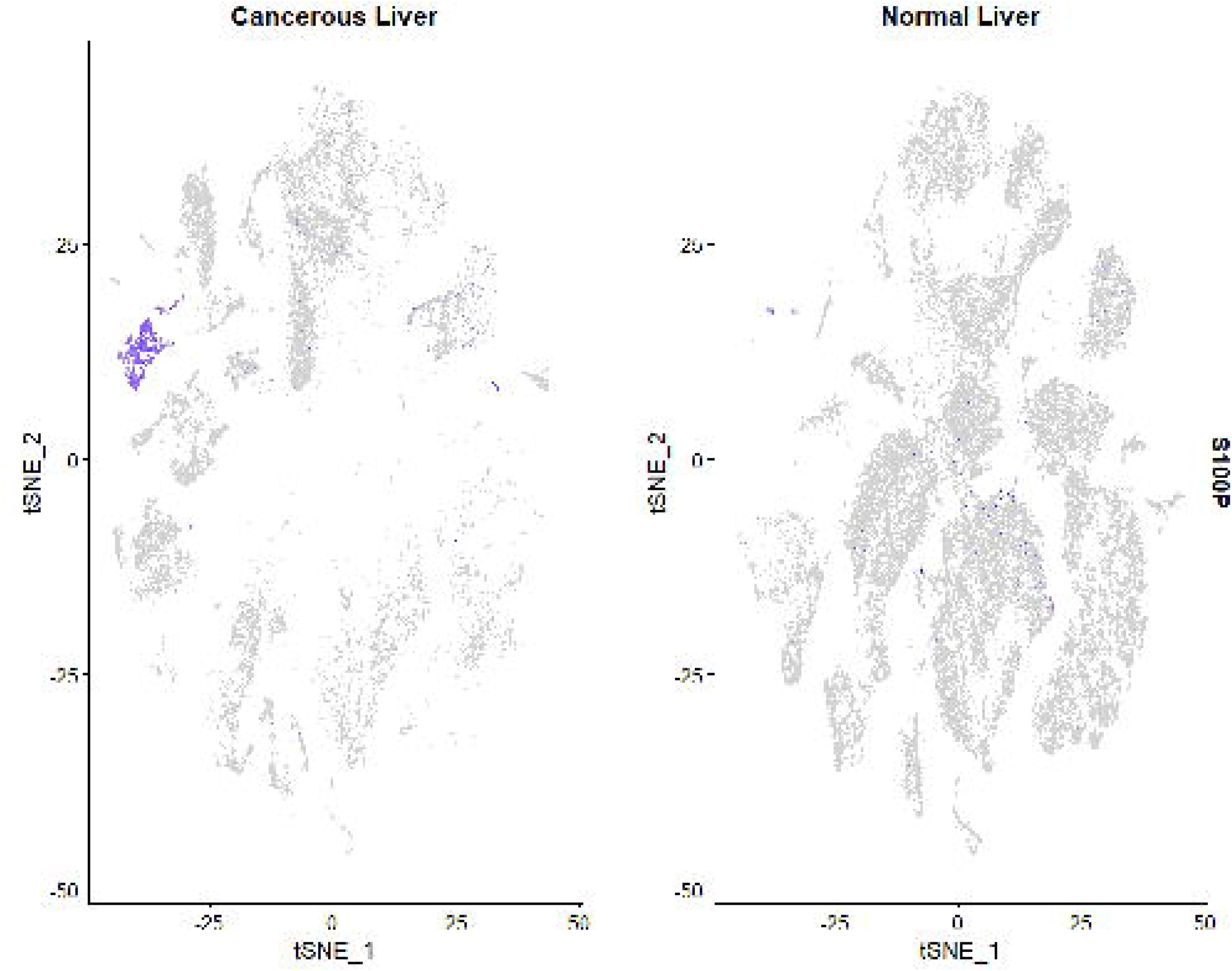

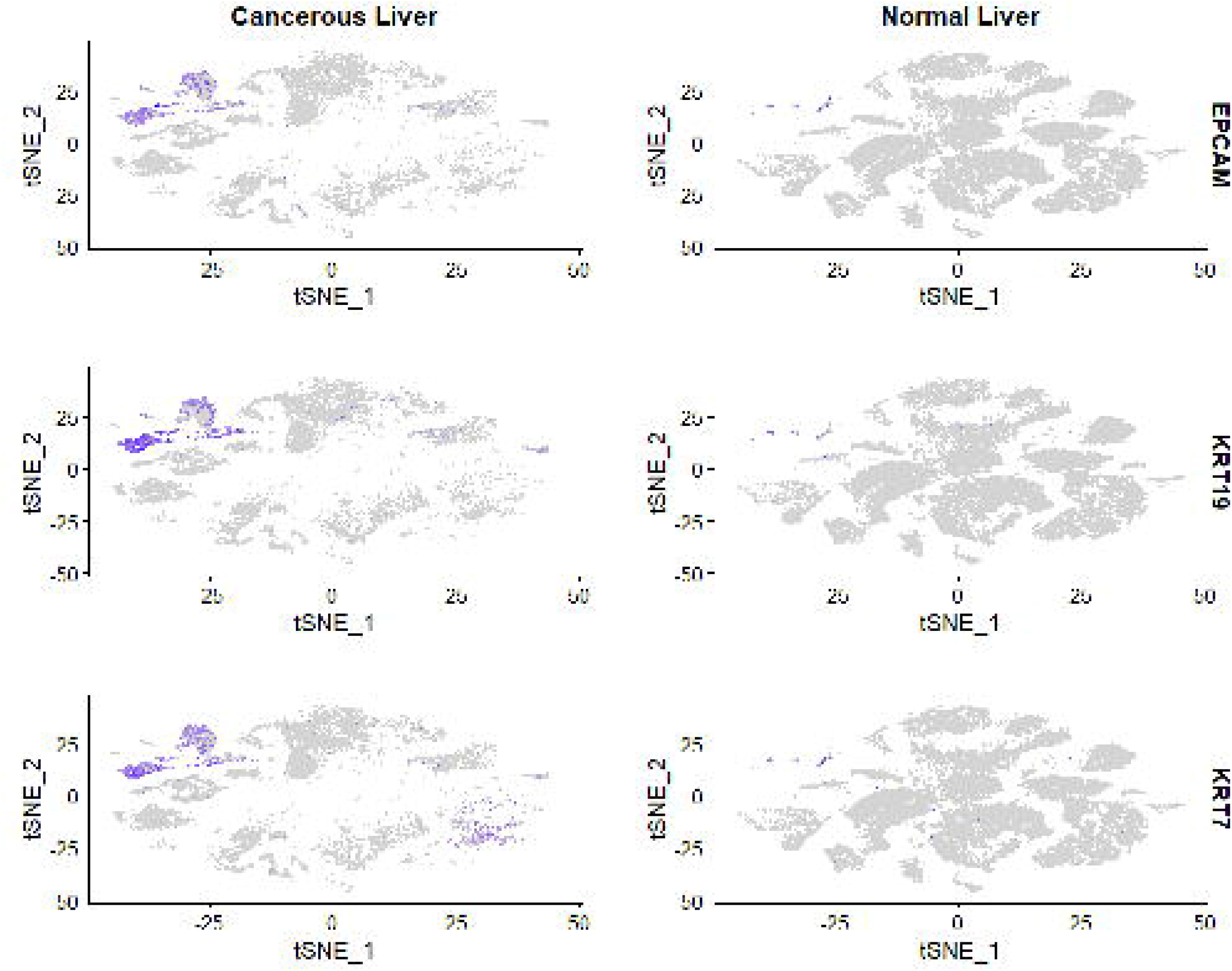

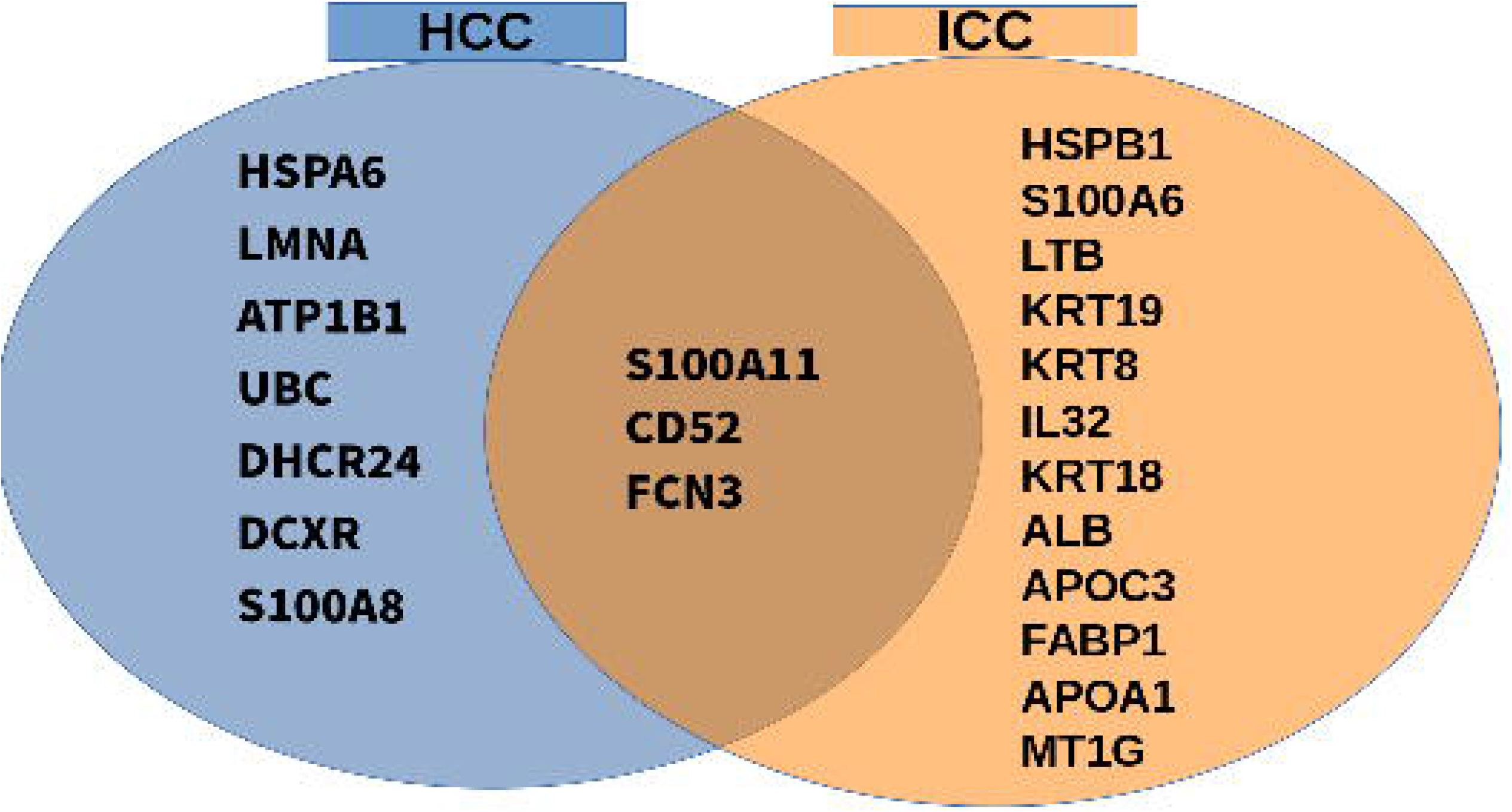

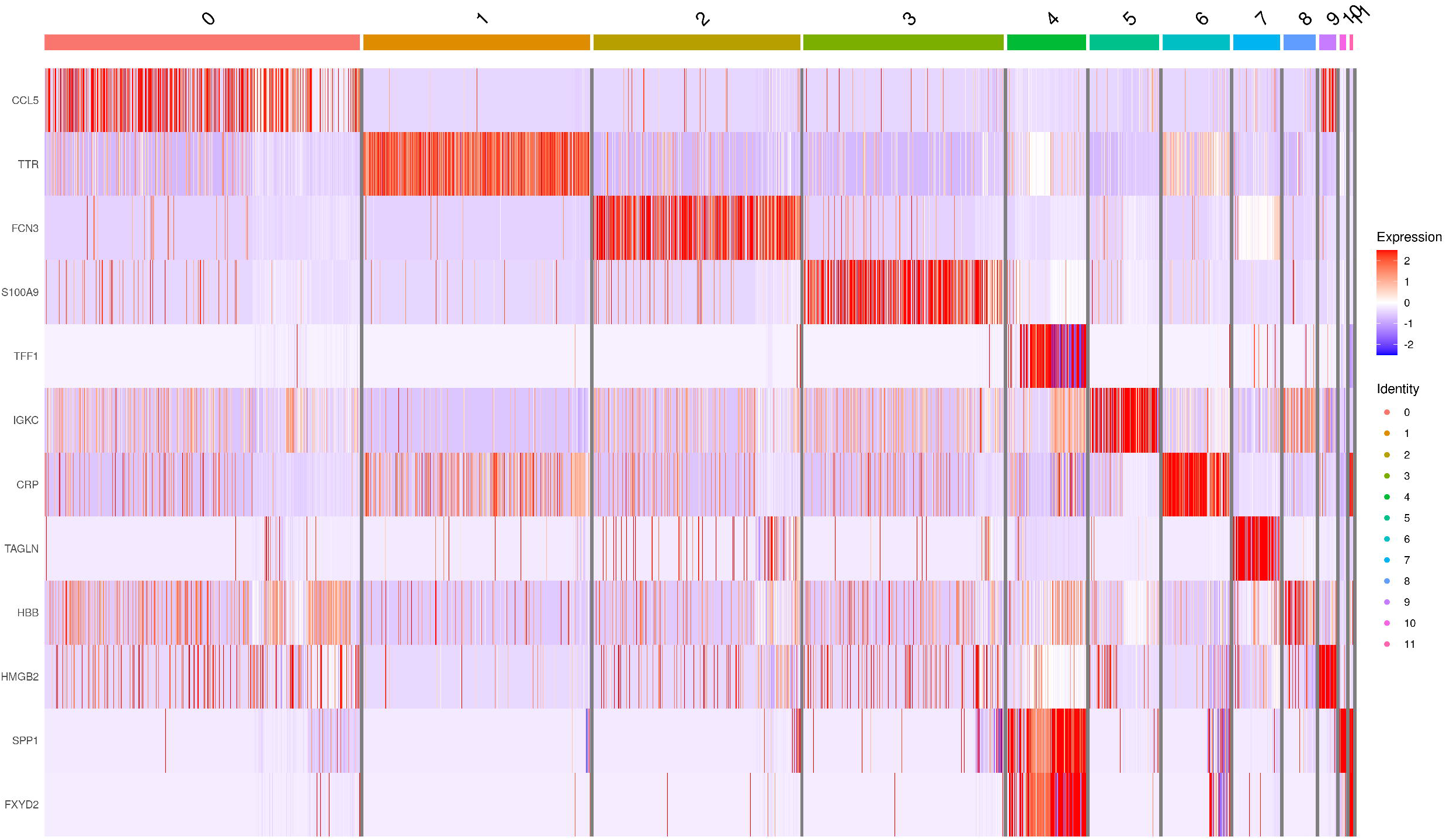

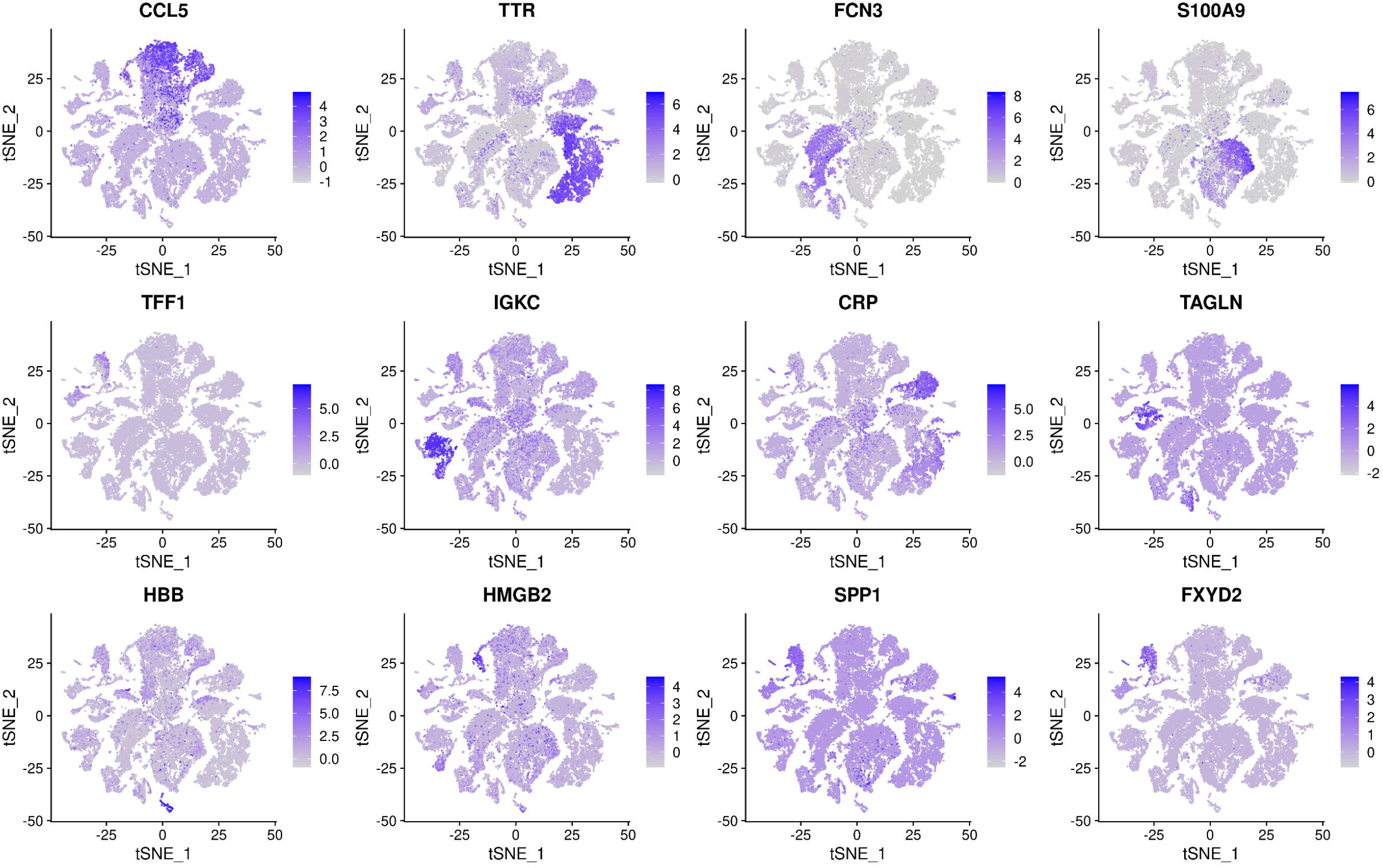

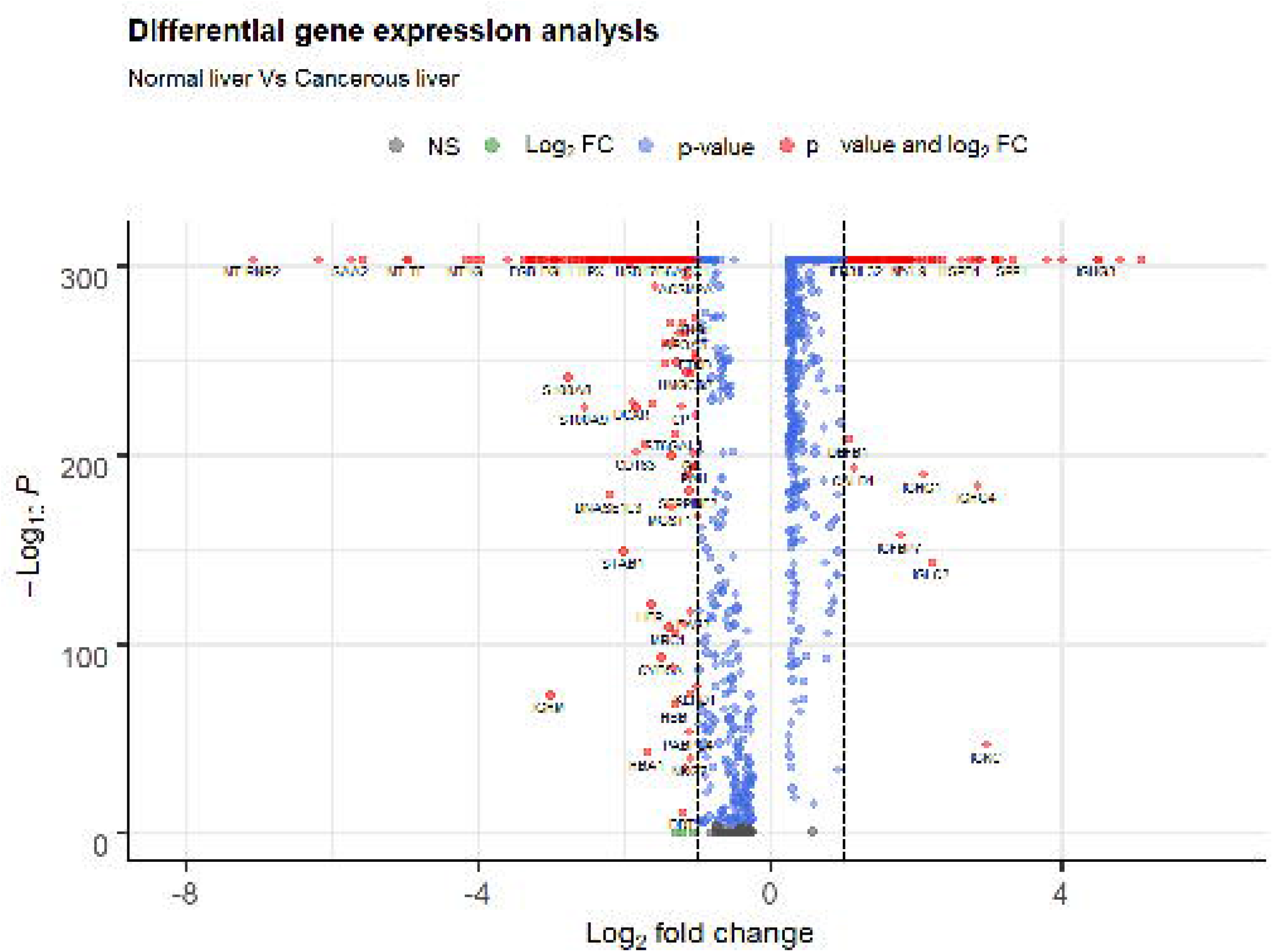

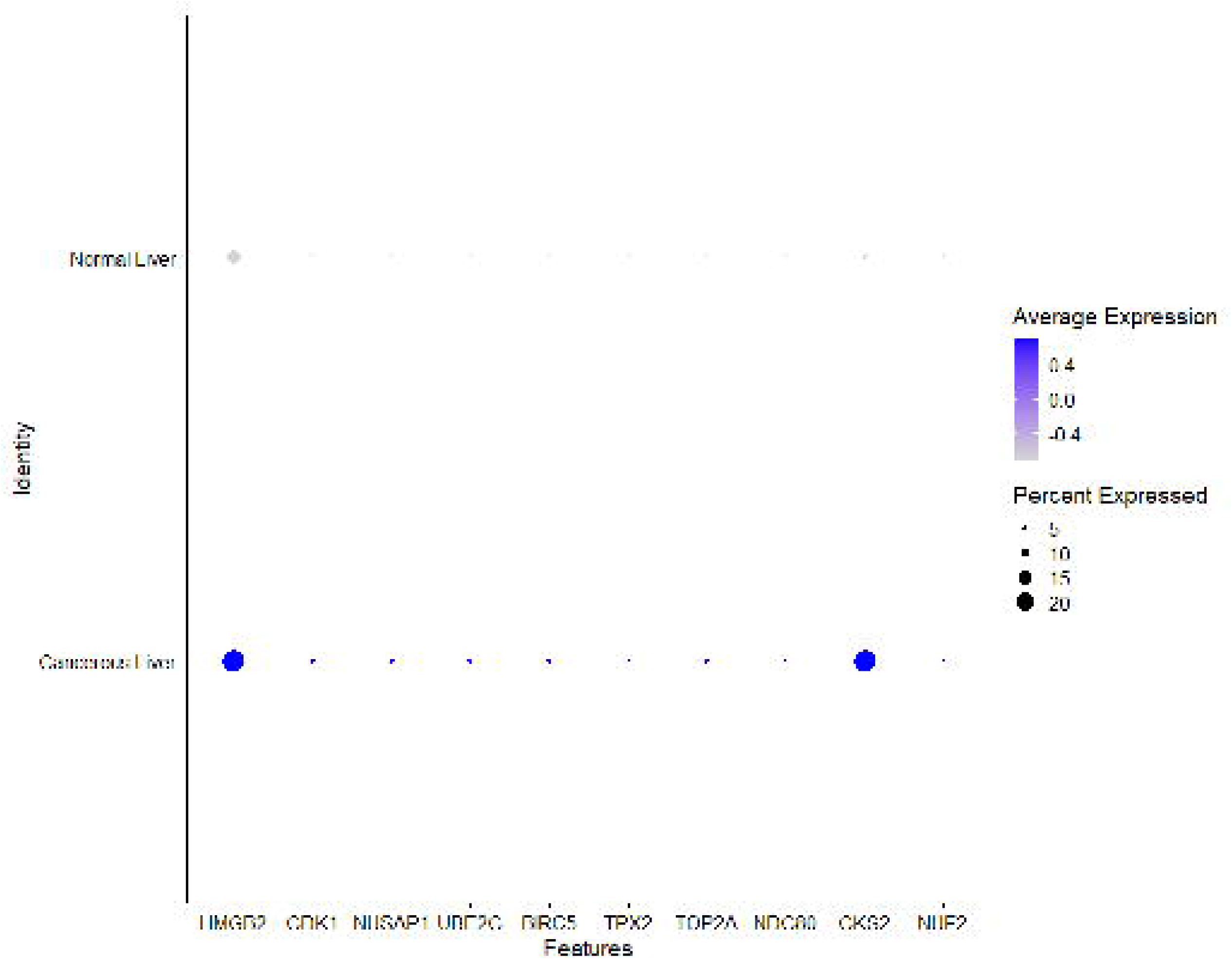

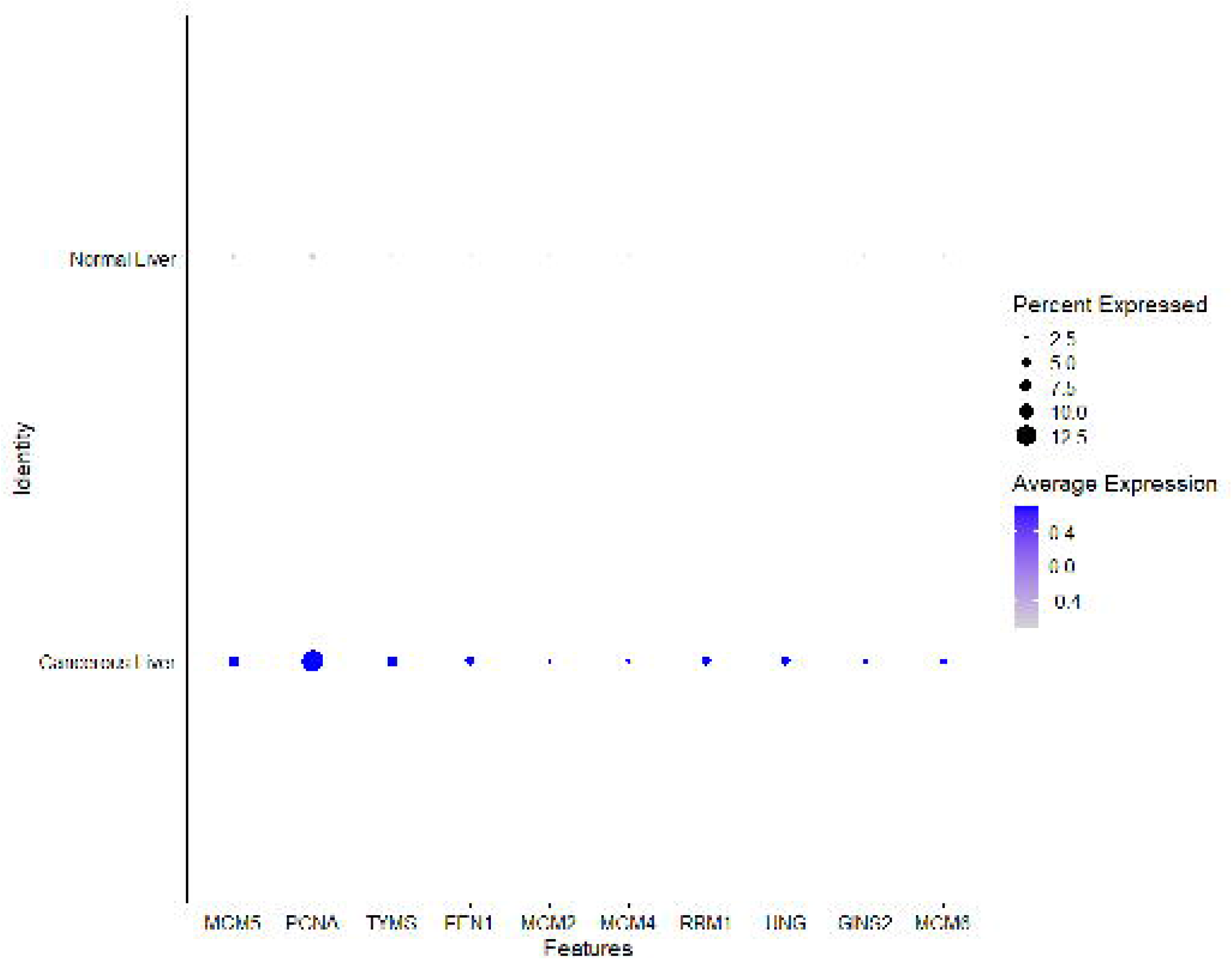

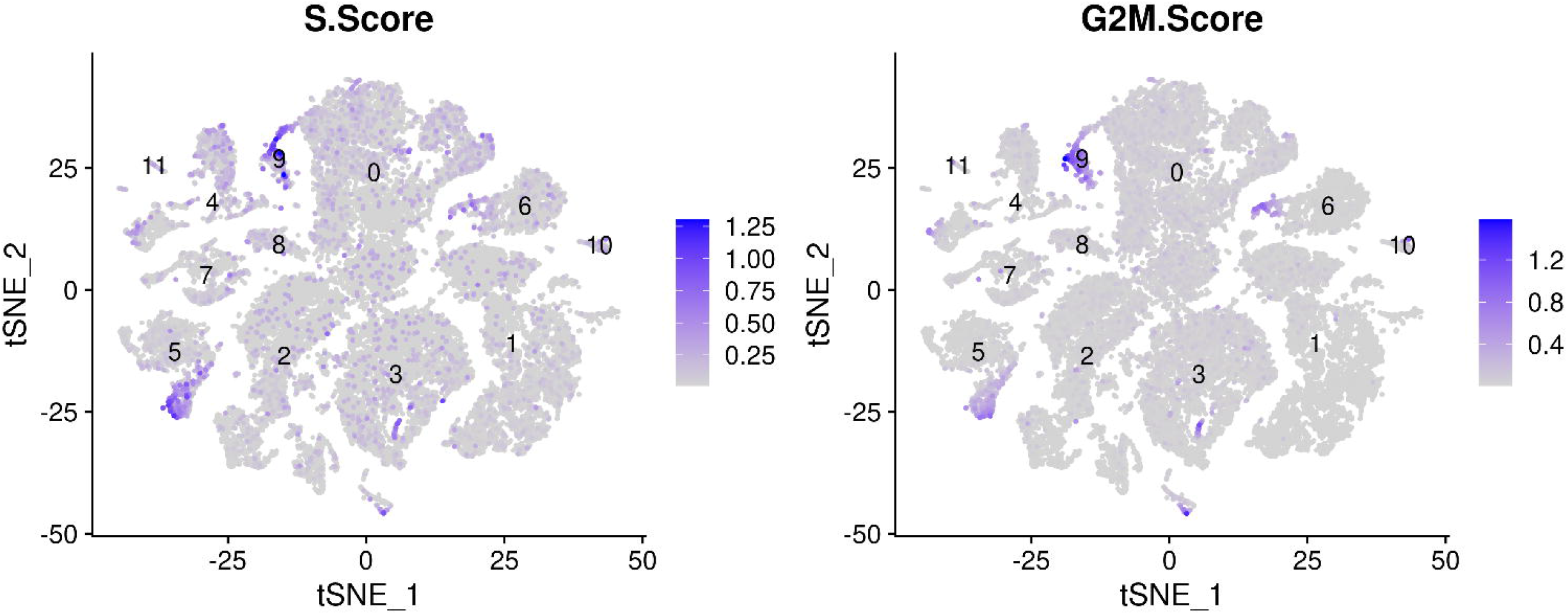

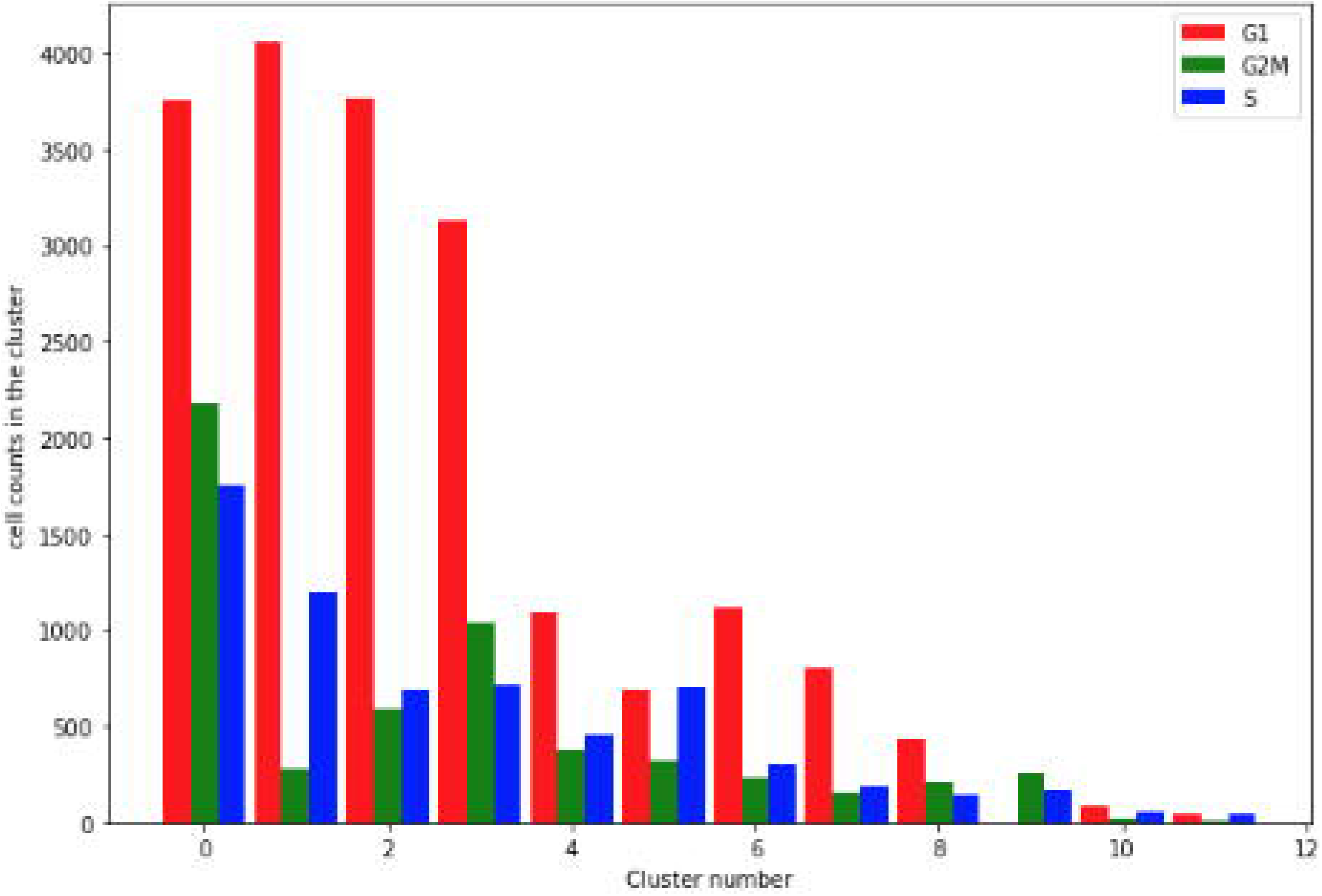

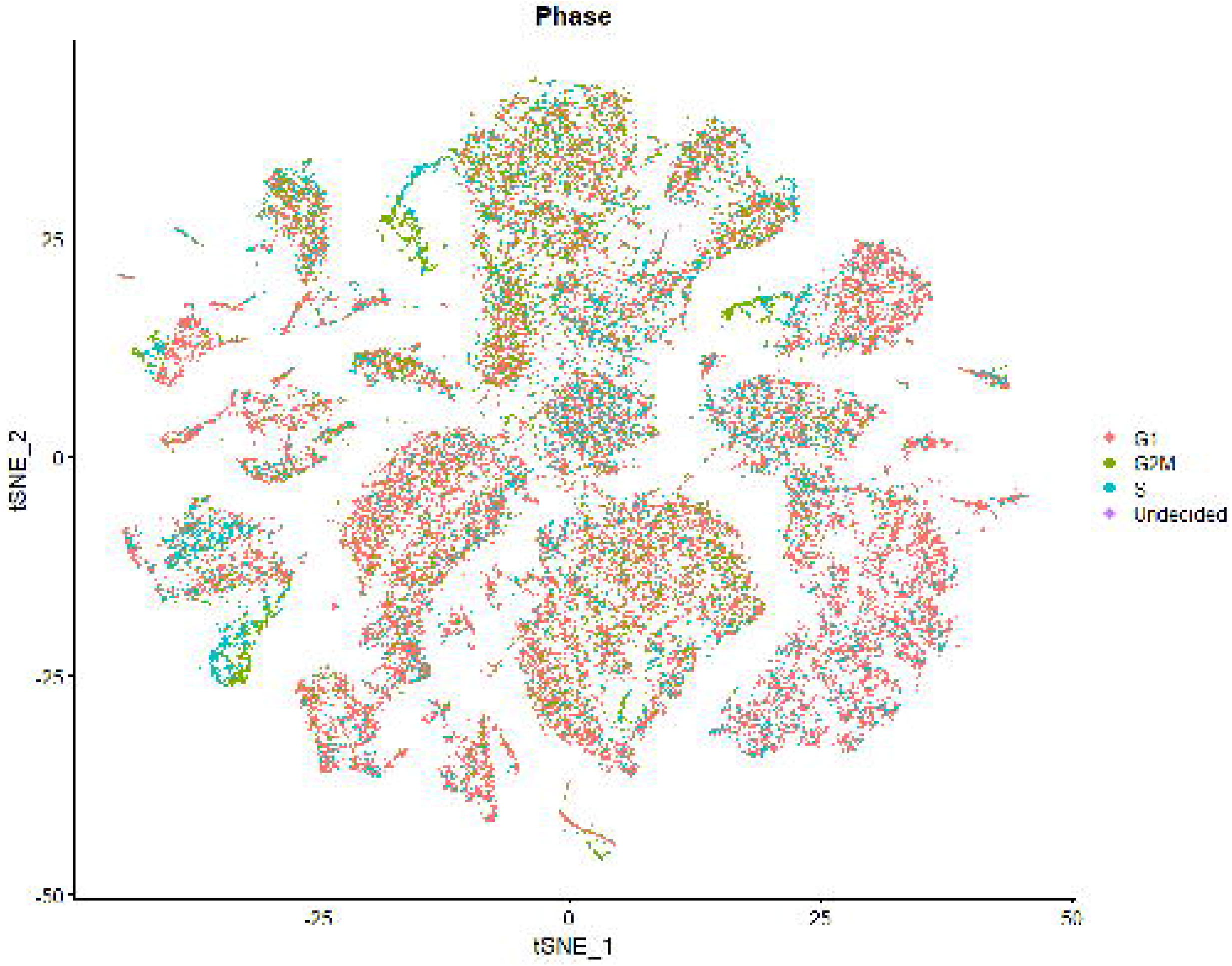

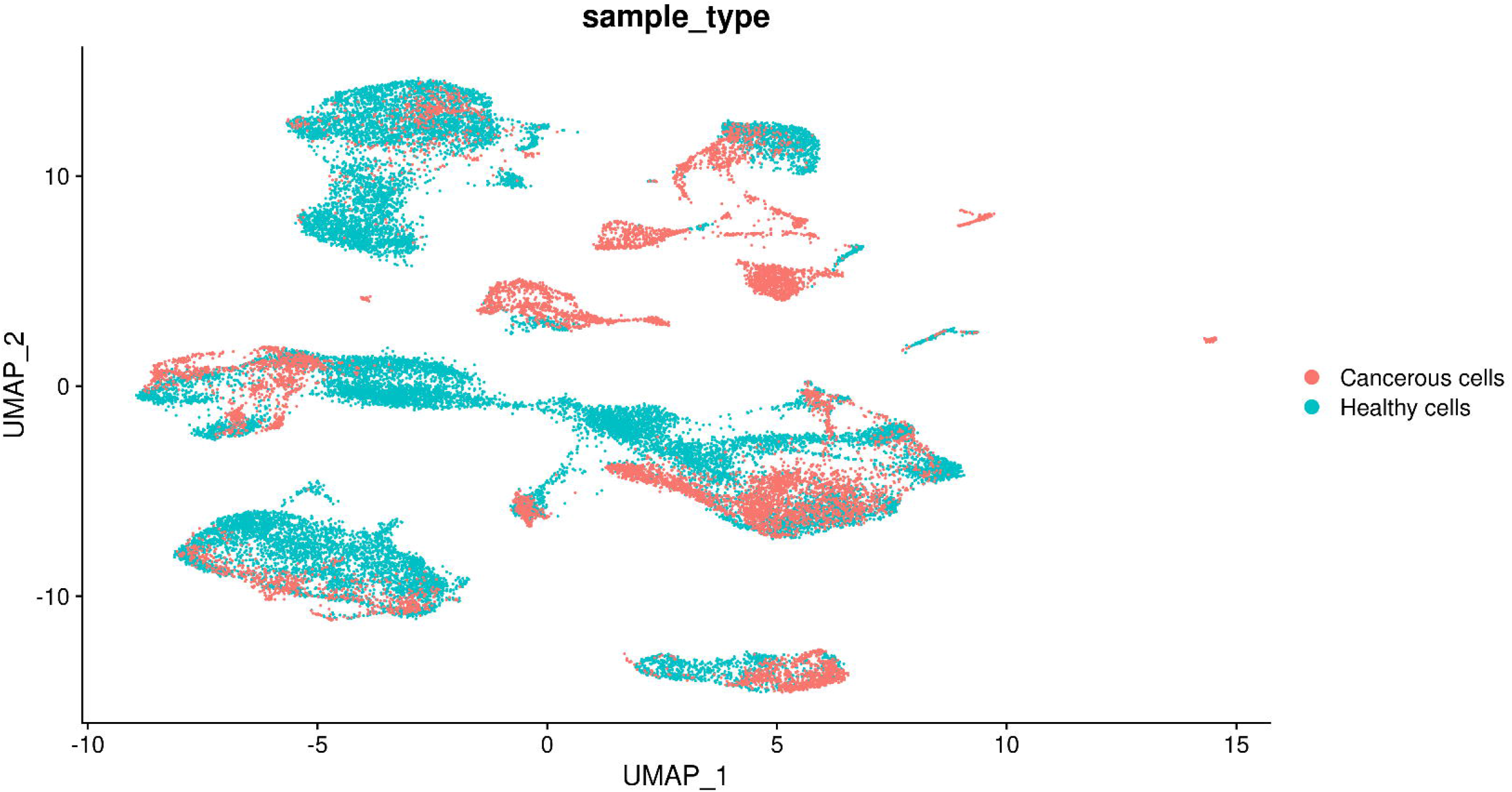

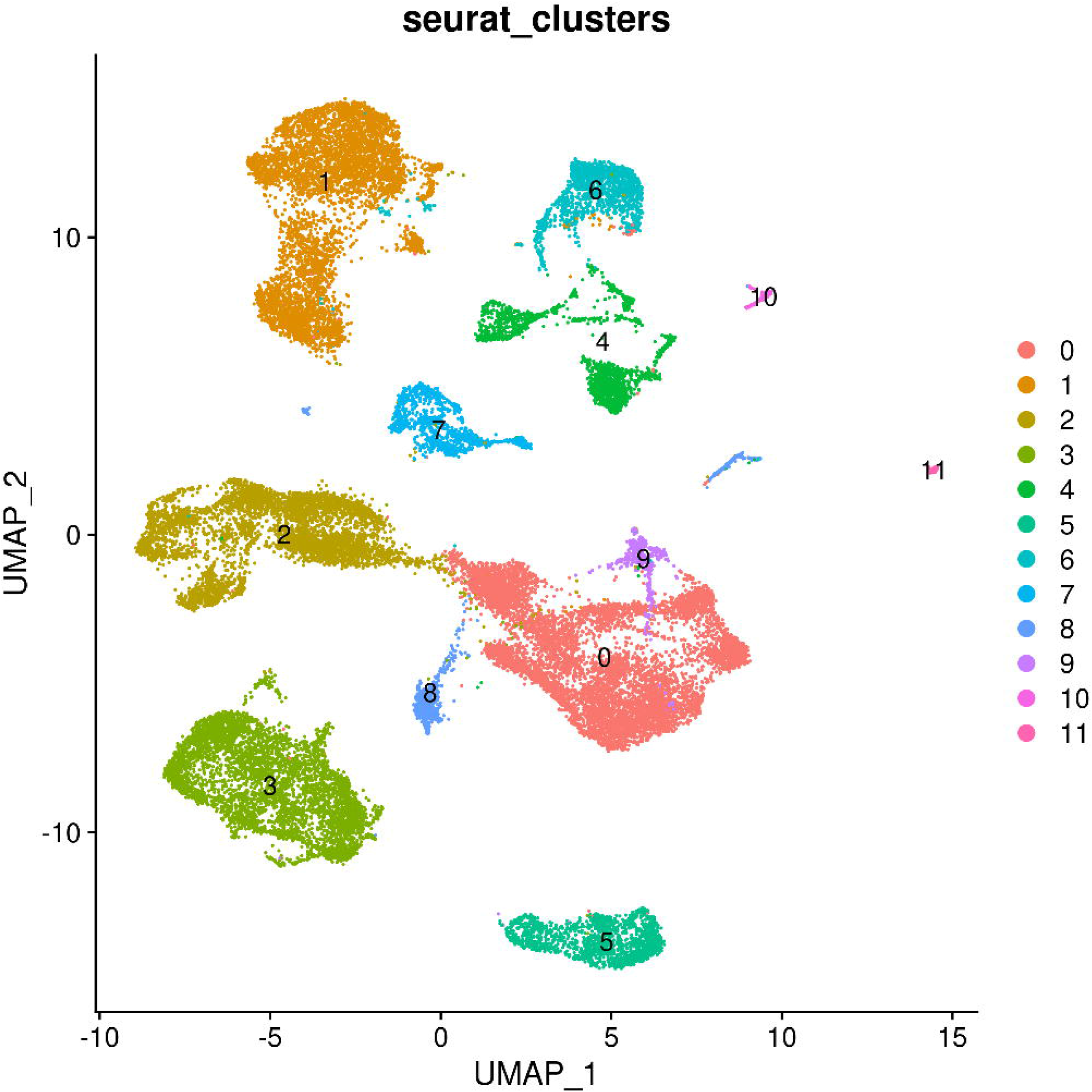

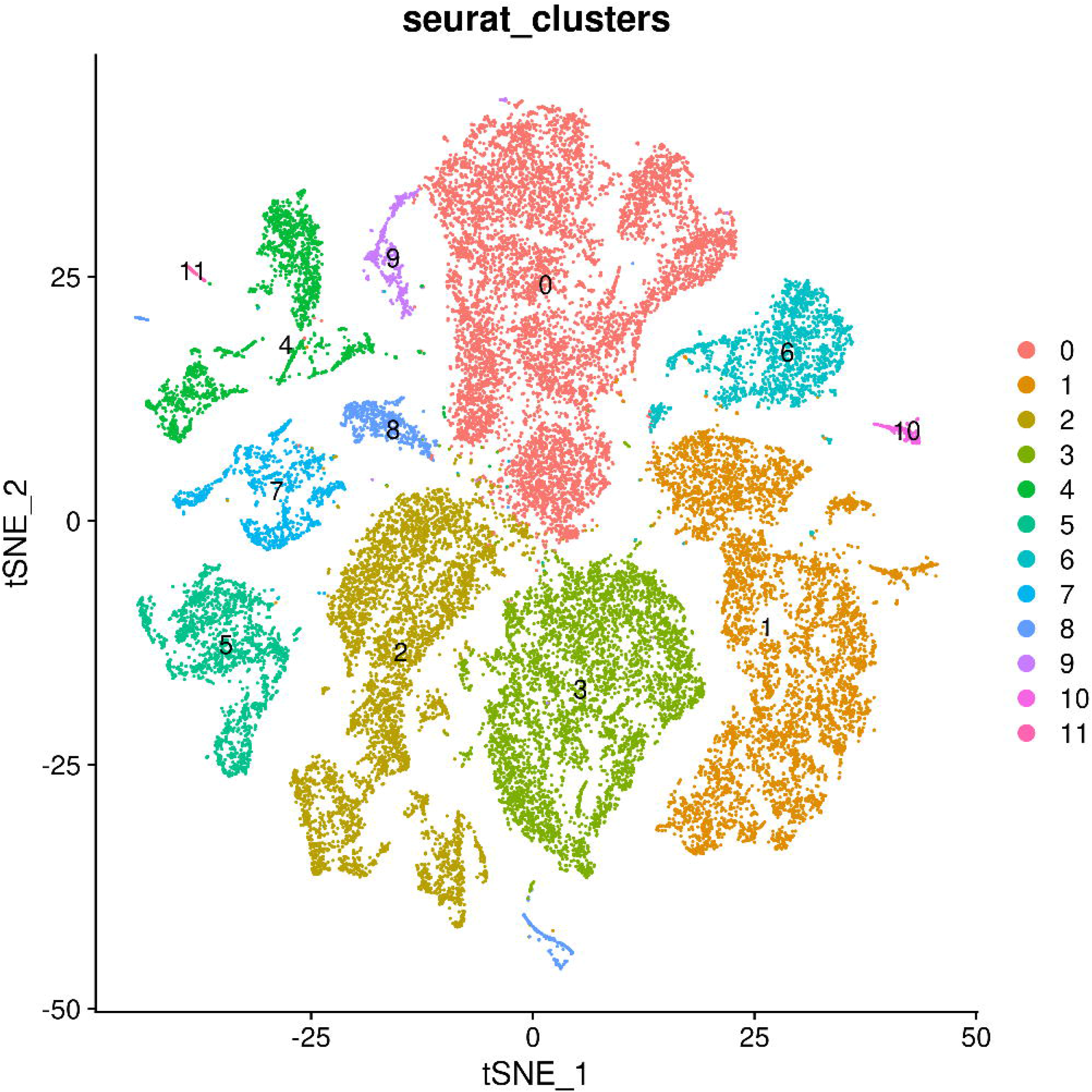

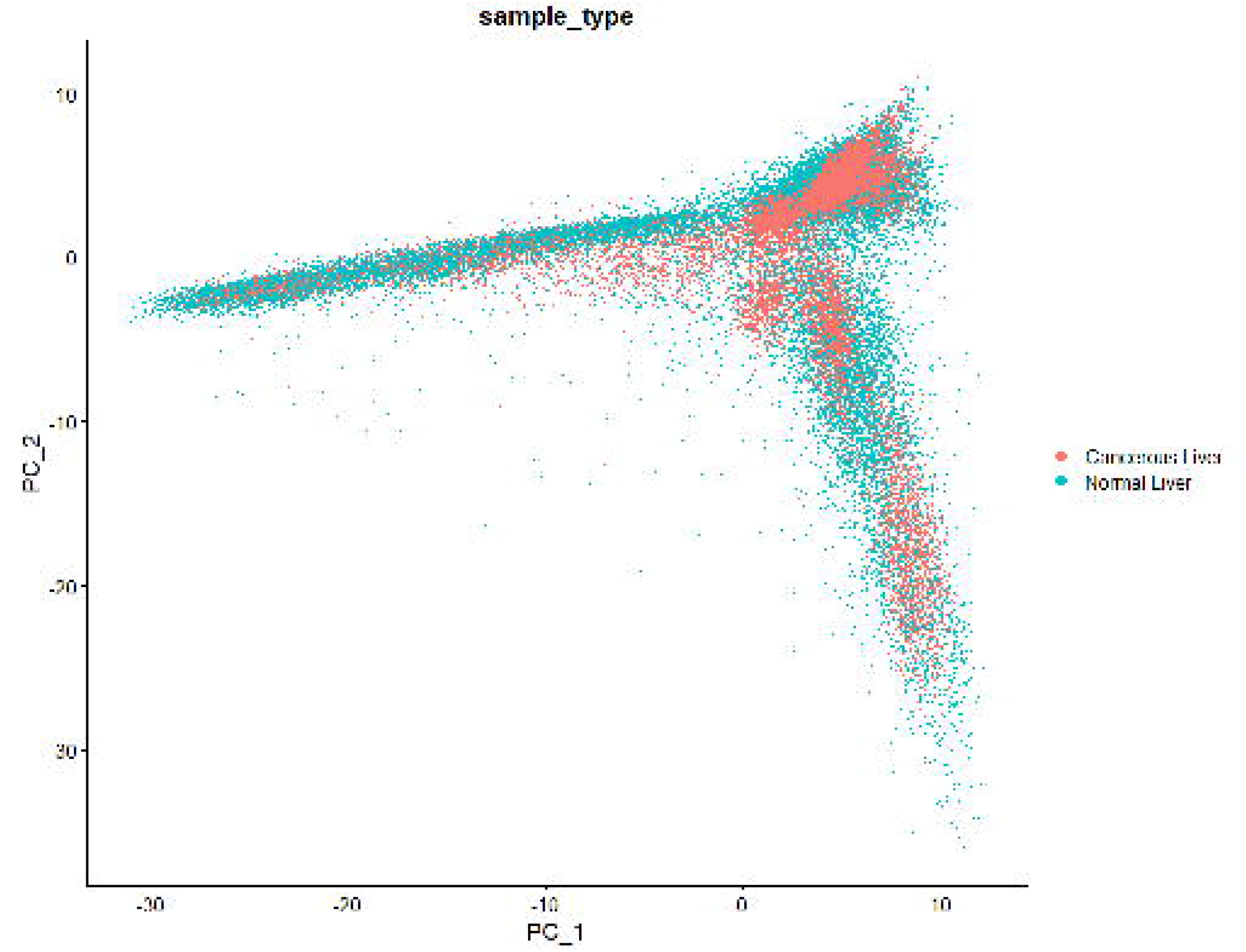

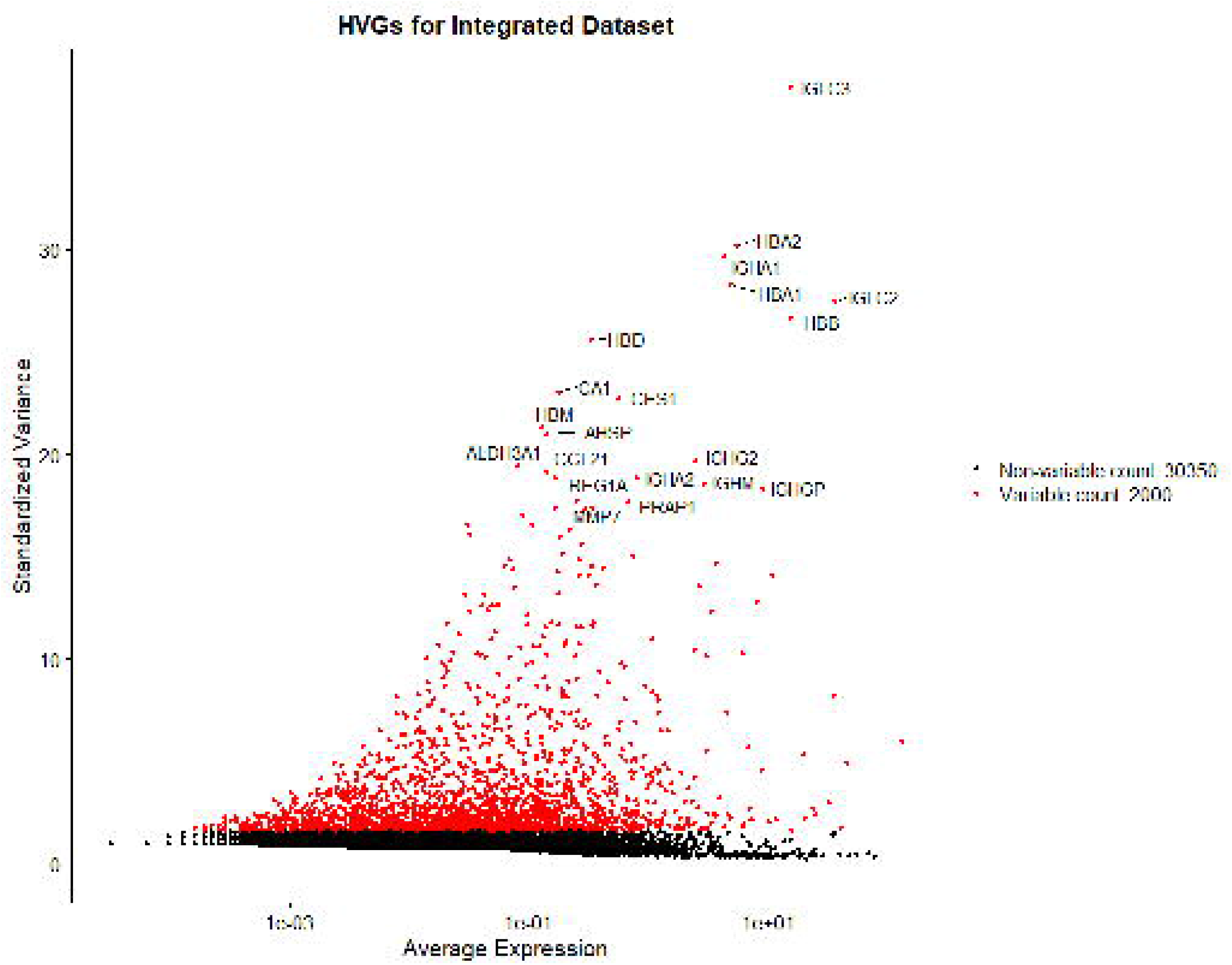

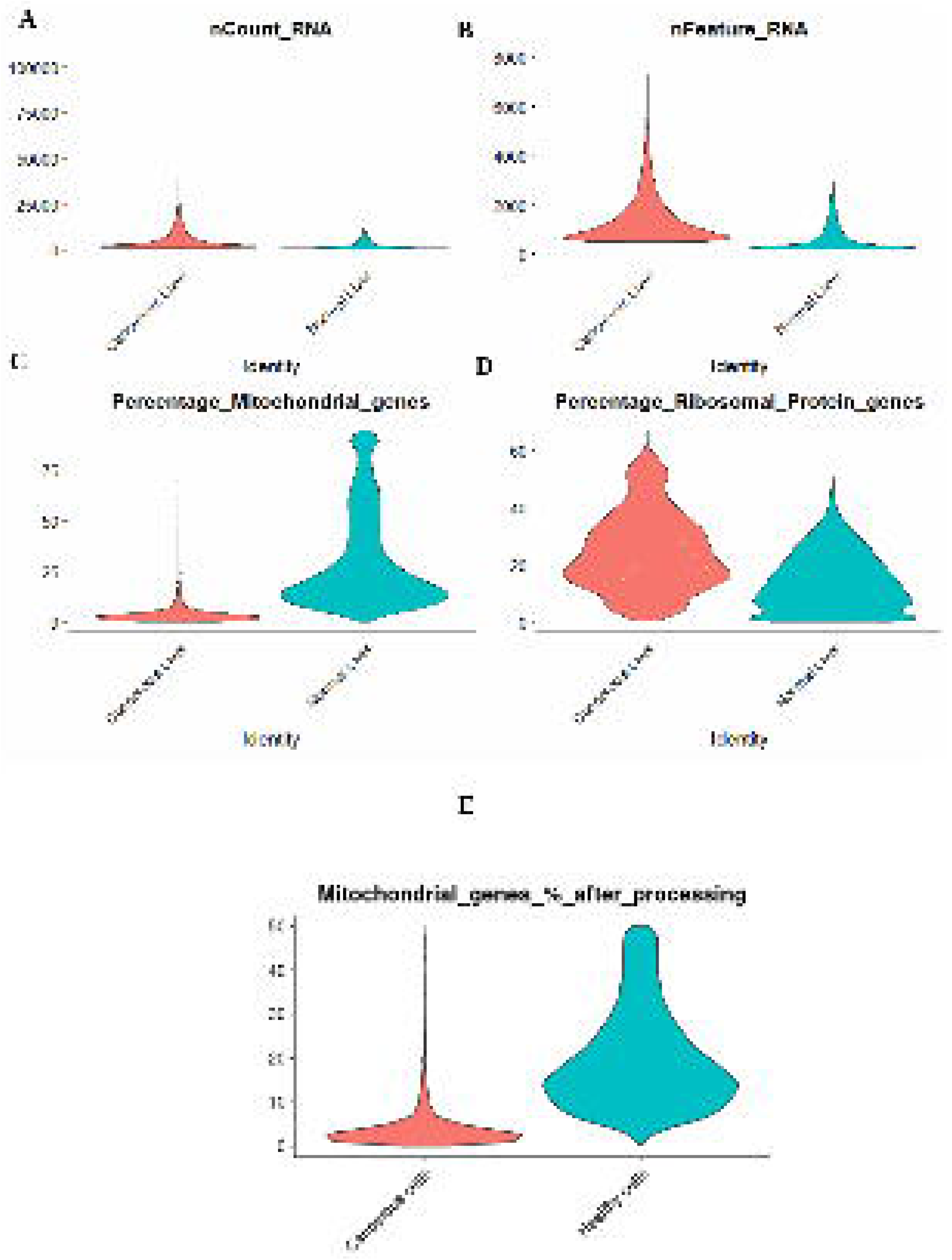

